# Azithromycin possesses biofilm–inhibitory activity and potentiates non-bactericidal colistin methanesulfonate against *Klebsiella pneumonia*

**DOI:** 10.1101/2022.02.17.480821

**Authors:** Olena V. Moshynets, Taras P. Baranovskyi, Scott Cameron, Olga S. Iungin, Ianina Pokholenko, Robyn Jerdan, Aleksandr Kamyshnyi, Alexey A. Krikunov, Viktoria V. Potochilova, Kateryna L. Rudnieva, Andrew J. Spiers

**Affiliations:** Institute of Molecular Biology and Genetics of the National Academy of Sciences of Ukraine, Kiev, Ukraine; Shupyk National Healthcare University of Ukraine, Kiev, Ukraine; Kyiv Regional Clinical Hospital, Kiev, Ukraine; School of Applied Sciences, Abertay University, Dundee, United Kingdom; Kyiv National University of Technologies and Design, Kiev, Ukraine; I. Horbachevsky Ternopil National Medical University, Ternopil, Ukraine; Amosov National Institute of Cardiovascular Surgery, Kiev, Ukraine

## Abstract

Novel antibiotic combinations may act synergistically to inhibit the growth of multidrug-resistant bacterial pathogens but predicting which will be successful is difficult, and standard antimicrobial susceptibility testing may not identify important physiological differences between planktonic free-swimming and biofilm-protected surface-attached sessile cells. Using a nominally macrolide-resistant model *Klebsiella pneumoniae* strain (ATCC 10031) we demonstrate the effectiveness of several macrolides in inhibiting biofilm growth in multi-well plates, and the ability of azithromycin (AZM) to improve the effectiveness of the antibacterial last-agent-of-choice for *K. pneumoniae* infections, colistin methanesulfonate (CMS), against biofilms. This synergistic action was also seen in biofilm tests of several *K. pneumoniae* hospital isolates and could also be identified in polymyxin B disc-diffusion assays on azithromycin plates. Our work highlights the complexity of antimicrobial-resistance in bacterial pathogens and the need to test antibiotics with biofilm models where potential synergies might provide new therapeutic opportunities not seen in liquid culture or colony-based assays.

## Introduction

Gram-negative infections are recognised as a serious global health problem with multi- and pan-drug resistant (M/PDR) strains included in the World Health Organisation’s list of antibiotic-resistant ‘priority pathogens’ that pose the greatest threat to human health. This includes *Klebsiella pneumoniae* which is a leading nosocomial causative agent of Gram-negative bacterial infections in Ukraine [1,2] and the frequency of M/PDR stains associated with a high mortality rate is increasing in eastern Europe and worldwide [3-5] and poses a significant challenge for the treatment of *K. penumoniae* infections.

An antibacterial last-agent-of-choice for M/PDR GNB strains is colistin methanesulfonate (CMS) (also known as colistimethate sodium and polymyxin E) which is an improved version of colistin sulfate [6,7]. CMS is generally regarded as a ‘weak antibiotic’ with a low bactericidal activity and clinical failure of 79% against colistin-sensitive strains at 14 days, and 43% mortality at 28 days [8]. However, CMS has the advantage of reduced kidney toxicity compared to colistin sulfate, allowing higher concentrations to be used and overcoming the poor bioavailability of the active form colistin (COL). Increasing CMS dosage can raise serum COL levels to ∼6 mg/L [9-12], but the minimum inhibitory concentration (MIC) breakpoint for COL-susceptible GNB is 2 mg/L and commonly used dosage CMS regimes have sub-optimal maximum serum concentration (C_max_) / MIC ratios. An alternative to increasing C_max_ is to use antibiotic combinations such as COL-carbapenem, COL-tigecycline and COL-aminoglycoside, which are more effective against carbapenemase-producing GNB [5,13] than monotherapy. The potentiation of weak antibiotics such as CMS with other antimicrobials may provide new therapeutic opportunities for the treatment of M/PDR Gram-negative bacterial infections.

However, not all antibiotic combinations make pharmaceutical sense and confusion may arise from predictions made of the efficacy of antimicrobials acting alone, and more combination tests are needed to identify potential synergistic activity in antibacterial combinations using physiologically relevant concentrations. Antimicrobial susceptibility tests such as the Kirby-Bauer disc-diffusion assay [14,15] which can be extended to identify synergistic actions between antibiotics [16-18] rely on colony growth across the surface of agar and may not accurately reflect the physiological and metabolic state of liquid culture (planktonic) bacterial cells or those encased in protective biofilms and adhered to surfaces. Biofilm-formation underlies the survival and persistence of many pathogens, providing phenotypic resistance or tolerance to therapy when genetic resistance might be restricted [19,20]. However, there are some reports of antimicrobials such as azithromycin (AZM) which are effective against *Pseudomonas aeruginosa* biofilms but not against colonies or liquid cultures [21-24], suggesting that new therapeutic opportunities for the treatment of M/PDR Gram-negative bacterial infections might also be found in re-testing antimicrobials using a biofilm model system rather than conventional agar plate or liquid-culture–based assays.

*K. pneumoniae* and *P. aeruginosa* share some common characteristics including the ability to form biofilms as well as similar macrolide resistance mechanisms [25]. We therefore hypothesized that AZM or other macrolides which have no effect against *K. pneumoniae* liquid cultures or colonies [26], might instead be effective against biofilm-forming *K. pneumoniae* cells (the ability to form biofilms is commonplace amongst *K. pneumoniae* isolates (e.g., 63 - 94% [27-29]) and MDR and biofilm-formation appear unlinked [29]). Moreover, we suggest that AZM or other macrolides acting as an anti-biofilm therapeutic agent might enhance the bactericidal effectiveness of CMS in a combined anti-*Klebsiella* therapy, as macrolides have been shown to potentiate CMS against both *K. pneumoniae* and *P. aeruginosa* [30]. Previously we have discussed the range of different aggregations bacteria form and key differences between classical biofilms, colonies, and liquid cultures [31]. In light of this and the recognition of the growing importance of biofilms in infections, the aims of our study were to evaluate the effectiveness of different macrolides in supressing biofilm growth by a model *K. pneumoniae* strain (ATCC 10031) [32-34] test the most effective macrolide suppressor in combination with CMS used in a range of physiologically-relevant concentrations, and then to test this combination using a collection of *K. pneumoniae* hospital isolates to evaluate the potential therapeutic benefit offered by a macrolide-CMS combination against *K. pneumoniae* biofilm– associated infections.

## Materials and methods

### *Klebsiella* strains

*Klebsiella pneumoniae* ATCC 10031 (Ukrainian National Collection ATCC) was used as a model strain for testing biofilm formation and free-living (planktonic) behaviour under different antibiotic treatments. Biofilm and planktonic responses to antibiotic treatment was also determined for eleven *K. pneumoniae* isolates from Ukrainian patients, including the multidrug (MDR) strains UHI 329, UHI 1090, UHI 1609, UHI 1633 and UHI 166, and the non-MDR strains UHI 117, UHI 486, UHI 489, UHI 509, UHI 519 and UHI 520 [32] (see **S1 Table** for strain origins and antibiotic sensitivities; strains were selected without knowing whether they could form biofilms or not). All strains were stored as frozen stocks in 25 (v/v) % glycerol at -80 °C.

### Culturing and antibiotics

*Klebsiella* were cultured aerobically at 37°C using Luria-Bertani (LB) medium (10 g/L peptone, 5 g/L yeast extract and 10 g/L sodium chloride; 12 g/L agar added for plates [35]) and Muller Hinton Agar (OXOID, UK). Strains were recovered from frozen stocks on LB plates before initiating over-night shaken cultures to provide fresh inocula for experiments (direct inoculation from frozen stocks was found to reduce siderophore production and biofilm development considerably; unpublished observations). Culture densities and dilutions were determined by OD_625_ measurements using a Spectronic Helios Epsilon spectrophotometer (Thermo Fisher Scientific, UK) with 10 mm optical-path cuvettes.

Eight macrolide antibiotics were used for the main study and included Azithromycin (AZM; Pharmex Group, Ukraine), Clarithromycin (CLR; Abbott, Italy), Erythromycin (ERM; Vitaminy, Ukraine), Josamycin (JSM; Astellas Pharma, Japan), Midecamycin (MID; KRKA, Slovenia), Roxithromycin (ROX; Darnitsa, Ukraine), Spiramycin (SPM; Sanofi Aventis, Italy), and Tylosin (TYL; Ukrzoovetprompostach, Ukraine). Stock solutions containing 9 g/L of each antibiotic were prepared using DMSO as solvent and kept at -20°C for no longer than 7 days before use. A 1 g/L stock solution of colistin methanesulfonate (CMS) (Forest Laboratories, UK) was prepared in water and used within 30 minutes of preparation. Discs containing antibiotics (HiMedia Laboratories, India) as well as AST-N332 cards for the VITEK 2 Advanced Expert System (bioMérieux, France) were used for antimicrobial susceptibility and sensitivity assays listed in **S1 Table**.

### Confirming antimicrobial susceptibilities of the UHI strains

The Kirby-Bauer disc-diffusion assay [14,15] was used to determine the antimicrobial susceptibility of the *K. pneumoniae* UHI strains. Mueller Hinton plates were spread with 200 µL of cell suspension prepared by diluting over-night cultures to 0.12 – 0.18 OD_625_ and allowed to dry before no more than five discs with different antibiotics per plate were applied. Susceptibility was then determined as the diameter (mm) of inhibition after incubation for two days and interpreted according to European Committee on Antimicrobial Susceptibility Testing (EUCAST) guidelines [26], HiMedia (for Cefoperazone/Sulbactam [36]), and the Ministry of Health of Ukraine (for Gatifloxacin [37]). β-lactamase genes were detected by PCR using DNA purified using AmpliSense DNA-sorb-B Nucleic Acid Extraction, MDR KPCOXA-48-FRT and MBL-FRT PCR kits (InterLabService, Russia). β-lactamase activity was determined with KPC&MBL&OXA-48 and ESBL+AmpC screen disc kits (Liofilchem, Italy) according to EUCAST guidelines [38]. AST-N332 cards were also used with the VITEK 2 Advanced Expert System to confirm the MDR strain phenotypes. Antimicrobial susceptibility testing of CMS was performed with the broth microdilution assay SensiTest™ Colistin (Liofilchem, Italy) results interpreted according to EUCAST guidelines [26].

The Kirby-Bauer disc-diffusion assay was also used to determine whether an AZM and polymyxin B synergistic action could be detected for UHI. Mueller Hinton plates supplemented with 0, 3, 6, 9 mg/L AZM were spread with 200 µL of cell suspension prepared by diluting over-night cultures to 0.12 – 0.18 OD_625_ and allowed to dry before replicate polymyxin-B PB300U discs (*n* = 3) were applied. Susceptibility was then determined as the diameter (mm) of inhibition after incubation for two days.

### Planktonic and biofilm growth assays

Planktonic growth assays of ATCC 10031 and the UHI strains was assessed in loosely lidded 30 ml glass vials which were incubated with shaking before assay. The vials contained 6 mL LB and AZM and CSM added to 6 mL LB vials to final concentrations of 0, 3, 6 and 9 mg/L AZM and 0, 2, 4, 6, 8, 12, 16, 18, 24, 28 and 32 mg/L CMS. Aliquots of over-night culture were added to replicate vials (*n* = 3) to ∼5 × 10^6^ CFU/mL and growth determined after 16 h incubation by removing 200 µL samples which were transferred to a polystyrene 96-well plate and OD_570_ measured using a BioTek ELx800 (BioTek Instruments, Winooski, VT, USA) plate reader.

Preliminary assessment of biofilm formation in vials and polystyrene 96-well plates found that ATCC 10031 and UHI solid-liquid (S-L) interface biofilms were very fragile with no growth occurring in the liquid column and no visible sediment from the initial inoculum. However, these biofilms could not be reliably retained during washing and Crystal violet staining following standard procedures (e.g., [29]). We therefor decided to modify our approach and used direct OD_570_ measurements of biofilms in situ in unprocessed 96-well plates. 20 µL aliquots of over-night culture were used to inoculate replicate wells (*n* = 8 for individual macrolide assays and *n* = 3 for AZM – CSM assays) containing 200 µL LB with the appropriate antibiotics (0, 1, 3, 6 and 9 mg/L macrolide for the for individual macrolide assays, 0, 3, 6 and 9 mg/L AZM and 0, 2, 4, 6, 8, 12, 16, 18, 24, 28 and 32 mg/L CMS for the AZM – CSM assays). Growth (OD_570_) was determined after 24 – 48 h static incubation using a BioTek ELx800 plate reader. Metabolic activity was also determined after adding MTT (3-(4,5-dimethylthiazol-2-yl)-2,5-diphenyltetrazolium bromide; Sigma-Aldrich, UK) to a final concentration of 0.05% (v/v) and a further incubation of 3 h (after [39]). Cells were recovered from each well by centrifugation at 6000 g for 15 minutes using an Eppendorf 5424 Microcentrifuge (Eppendorf, Hamburg, Germany). The cell pellet was dissolved in 250 µL of DMSO, 125 µL of which was transferred to a polystyrene 96-well plate and metabolic activity (A_570_) measured using a BioTek ELx800 plate reader. We consistently found greater variation in A_570_ measurements of biofilm growth than in OD_570_ measurements which suggest that the MTT assay was the technically more difficult assay to perform.

### Statistical analyses

Experiments were performed with replicates (*n*) and data examined using JMP v12 statistical software (SAS Institute Inc.). Means ± standard errors are shown, and degrees of freedom (DF), test statistics, alpha and p-values provided. A mixed-effects modelling approach was used to investigate the effect of treatments on relative planktonic and biofilm growth and colony inhibition. The Restricted Maximum Likelihood (RML) method with unbounded variance components was used with experimental replicate included as a random effect in all models, and antibiotics, antibiotic concentrations, strain type (MDR and non-MDR), and strain nested in type (strain[type]), as fixed effects. Models were run iteratively with the residuals inspected and outliers removed until normality was attained as determined by Shapiro-Wilks tests (p < 0.05). Model robustness was monitored by RSquare and the retention of degrees of freedom with concentration ranges limited to avoid singularities. Effects and interactions were investigated using Least-Squares Means (LSMeans) Differences Student’s and Tukey HSD tests (alpha = 0.05).

For clarity, individual mixed-effects models were run with antibiotic concentration as an effect to differentiate treatments and the results of the LSMeans Tukey HSD tests are shown in the figures. Spearman’s (rho) coefficient was used to test for correlations between mean OD_570_ and A_570_ data. Some difficulty was encountered in analysing the relative inhibition data for individual strains using a mixed-effects modelling approach where normality could not be achieved (some data were too discrete) and one strain was omitted from the combined mixed-effects model as there was no variation in relative inhibition across AZM concentrations. For some, a general linear modelling (GLM) approach was successful with replicate and antibiotic concentration included as effects. However, two strains could not be analysed using GLMs or one-way ANOVA, so differences between key treatments were investigated with Wilcoxon (Rank sums) tests using a one-way test Chi-Square approximation. Rho was used to test for correlations between relative inhibition diameter and antibiotic concentration. Similarities in the effects of macrolides and the response of strains to antibiotics were visualised by Hierarchical Cluster Analysis using the Ward method and with equal weightings of factors. A Chi-Square test was used to test the independence of HCA clusters and MDR and non-MDR strain types (iCalcu.com).

## Results and discussion

### Biofilm growth by a model *Klebsiella* and suppression by macrolide antibiotics

*K. pneumonia* ATCC 10031 was found to form solid-liquid (S-L) interface biofilms [31] at the bottom of static LB microcosms within 24 h as had been reported earlier under different culture conditions [32-34]. As the liquid columns remained transparent with bacteria localised in the biofilm (*K. pneumoniae* is non-motile [40]), we used multi-well plates in which optical density measurements are made vertically up through the plate bottom and liquid column to record biofilm growth. This had the advantage of being able to quantify very weak biofilms that might not be retained during washing and elution of Crystal violet as noted in other cases [41] (ATCC 10031 produces weak biofilms compared to those produced by *Pseudomonas aeruginosa* and *Staphylococcus aureus* [33] or *Bordetella avium* [42]). We used this approach with *K. pneumonia* ATCC 10031 to evaluate the effect of eight macrolides, AZM, CLR, ERM, JSM, MID, ROX, SPM and TYL. These were tested over a concentration range of 0 – 9 mg/L and relative biofilm growth (OD_570_) determined after 24 h in comparison with a no-antibiotic control (**Fig. 1**). Of the eight macrolides, AZM and JSM showed high levels of inhibition of 2.5 – 5x with 3 – 9 mg/L of each antibiotic. CLR, ERM, MID and SPM showed moderate levels of inhibition of 1.2 – 1.7x over the same range, while ROX and TYL had little or no significant impact under the conditions tested here (LSMeans Differences Tukey HSD, alpha = 0.05 for individual macrolide mixed-effects models of relative OD_570_). Further analysis of all macrolides in a combined mixed-effects model of relative OD_570_ confirmed macrolide-type and concentration effects (*macrolide, F*_7,7_ = 126, p = 0.01; *concentration, F*_3,3_ = 290, p = 0.01; no-macrolide control omitted to avoid singularities), as well as a significant synergistic interaction between macrolides and macrolide concentration (*macrolide x concentration, F*_21,21_ = 13, p = 0.01). Similar results were found for biofilm assays incubated for 72 h (data not shown), and these findings demonstrate that *K. pneumonia* ATCC 10031 biofilm growth is differentially affected by both the type of macrolide and the concentrations tested in this assay.

**Figure 1.**
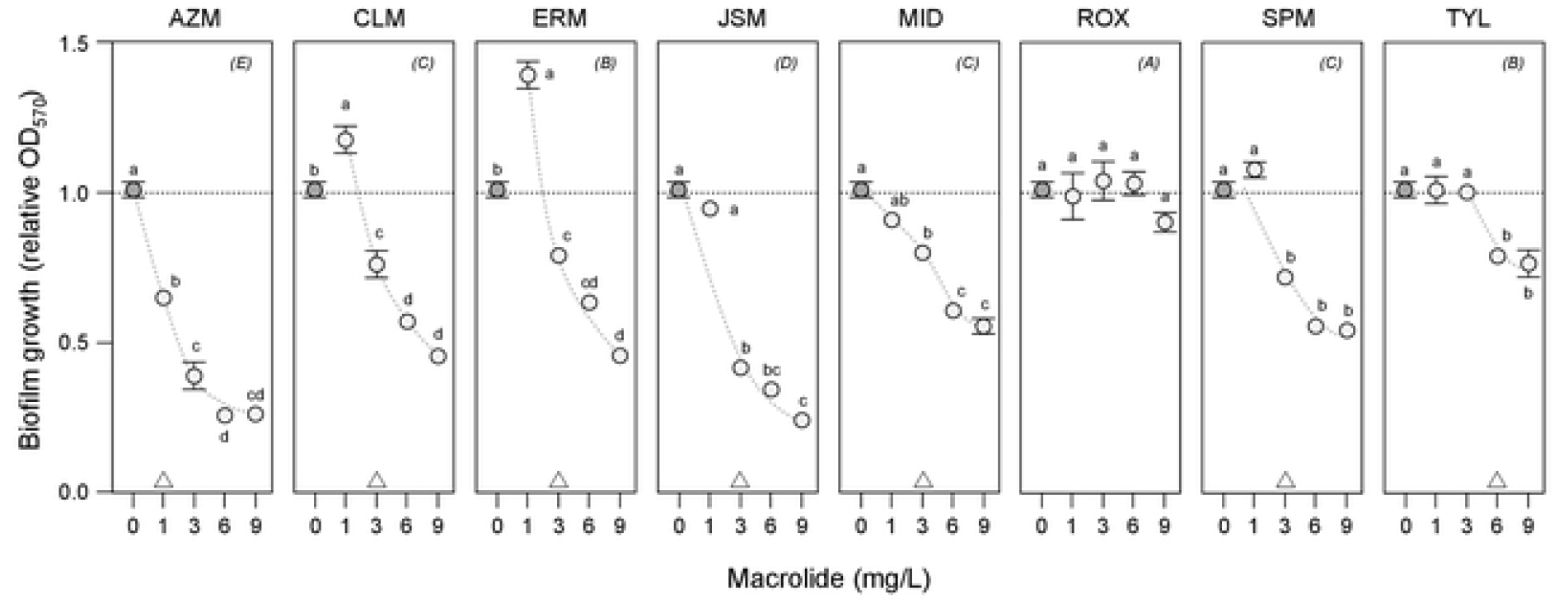
Optical density measurements of *K. pneumonia* biofilm growth show inhibition by macrolides. The inhibition of *K. pneumonia* ATCC 10031 biofilm growth by eight macrolides was investigated by biofilm assays with optical density (OD_570_) measurements after 24 h incubation. Mean relative OD_570_ ± SE (*n* = 3) is shown, with the no-macrolide (control) treatment indicated by dark circles and 1 – 9 mg/L macrolide treatments by light circles (trends are suggested by dashed curves). Means not connected by the same letters within panels are significantly different (LSMeans Differences Tukey HSD, alpha = 0.05 for individual macrolide mixed-effects models; note no significant differences for ROX). Triangles at the bottom of panels indicate biofilm minimum inhibitory concentrations (MIC) where relative OD_570_ was first significantly reduced by macrolide treatment. Macrolides not connected by the same letters indicated in parentheses in the top-right of each panel are significantly different (LSMeans Differences Tukey HSD, alpha = 0.05 for combined macrolide mixed-effects model of relative OD_570_).

These findings were confirmed using MTT and relative absorbance measurements (A_570_) to determine metabolic activity as a measure of biofilm growth rather than optical density–based measurements (**Fig. 2**). Significant levels of inhibition were seen for AZM and JSM and no effect for ROX and TYL (LSMeans Differences Tukey HSD, alpha = 0.05 for individual macrolide mixed-effects models of relative A_570_). Macrolide-type, concentration effects and the synergistic interaction were confirmed in a combined mixed-effects model of relative A_570_ (*macrolide, F*_7,7_ = 20, p = 0.01; *concentration, F*_3,3_ = 41, p = 0.01; *macrolide x concentration, F*_21,21_ = 2, p = 0.03; no-macrolide control omitted to be consistent with the relative OD_570_ model). Relative A_570_ treatment means were positively correlated with the relative OD_570_ means of the first assay (rho = 0.80, p = 0.01), demonstrating that the optical density and metabolic measurements of biofilm growth were in agreement. Hierarchical cluster analysis using relative A_570_ and OD_570_ means also grouped macrolides according to inhibition levels, with AZM and JSM clustered together and distal to both ROX and TYL (see **S1 Fig**. for the HCA dendrogram). As AZM is available in both peroral and parenteral forms for treatment, it was selected for further testing with CMS as a combined therapy against biofilm associated *Klebsiella* infections.

**Figure 2.**
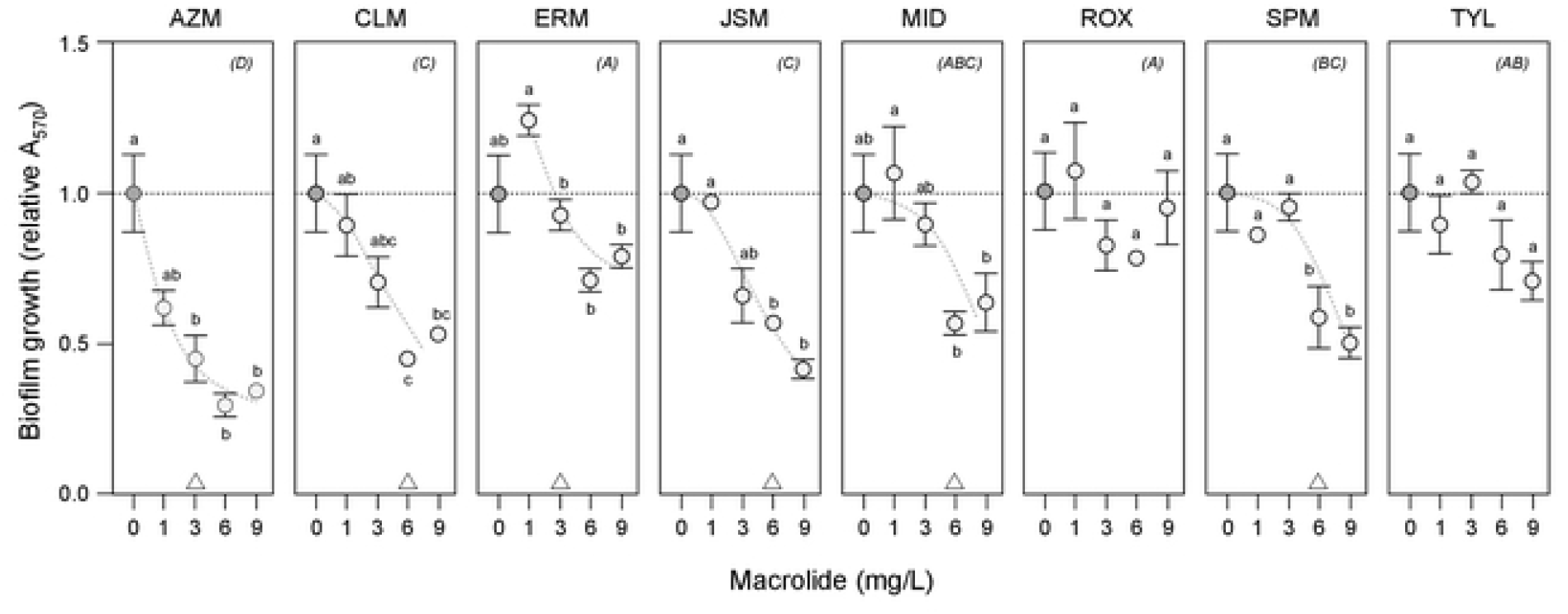
MTT assays of *K. pneumonia* biofilm growth show inhibition by macrolides. The inhibition of *K. pneumonia* ATCC 10031 biofilm growth by eight macrolides was investigated by biofilm assays using the metabolic indicator dye MTT with absorbance (A_570_) measurements after 24 h incubation. Mean relative A_570_ ± SE (*n* = 3) is shown, with the no-macrolide (control) treatment indicated by dark circles and 1 – 9 mg/L macrolide treatments by light circles (trends are suggested by dashed curves). Means not connected by the same letters within panels are significantly different (LSMeans Differences Tukey HSD, alpha = 0.05 for individual macrolide mixed-effects models). Triangles at the bottom of panels indicate biofilm minimum inhibitory concentrations (MIC) where relative A_570_ was first significantly reduced by macrolide treatment. Macrolides not connected by the same letters indicated in parentheses in the top-right of each panel are significantly different (LSMeans Differences Tukey HSD, alpha = 0.05 for combined macrolide mixed-effects model of relative OD_570_; note no significant differences for ROX and TYL).

### Antibacterial activity of AZM and CMS in combination against planktonic and biofilm–forming ATCC 10031 cells

The synergistic action of AZM and CMS on *K. pneumonia* ATCC 10031 was investigated using both planktonic and biofilm assays, because we hypothesized that lifestyle differences affect antibiotic sensitivities, and because infections involve both lifestyles [43,44]. CMS had no impact on planktonic growth when tested alone but inhibited growth at 6 mg/L with 3 mg/L AZM and at 2 mg/L with 9 mg/L AZM (**Fig. 3A**). A significant synergistic interaction between AZM and CMS was found in a mixed-effects model of relative OD_570_ (*AZM x CMS, F*_12,12_ = 170, p = 0.01) confirming the synergistic action of AZM and CMS on planktonic growth, as well as significant concentration effects (*AZM, F*_2,2_ = 3464, p = 0.01; *CMS, F*_6,6_ = 314, p = 0.01; data limited to 0 – 16 mg/L CMS to avoid singularities as higher-concentration treatments were highly correlated). In comparison, CMS was found to have an inhibitive effect on biofilm growth without AZM and inhibited growth at a lower concentration (2 mg/L) with 3 mg/L AZM (**Fig. 3B**) (growth of the initial inoculum in these assays was fully suppressed by 9 mg/L AZM). Once again, a significant synergistic interaction was found in a mixed-effects model of relative OD_560_ (*AZM x CMS, F*_12,12_ = 176, p = 0.01) as well as significant concentration effects (*AZM, F*_2,2_ = 1786, p = 0.01; *CMS, F*_6,6_ = 354, p = 0.01; data limited to be consistent with the planktonic model).

**Figure 3.**
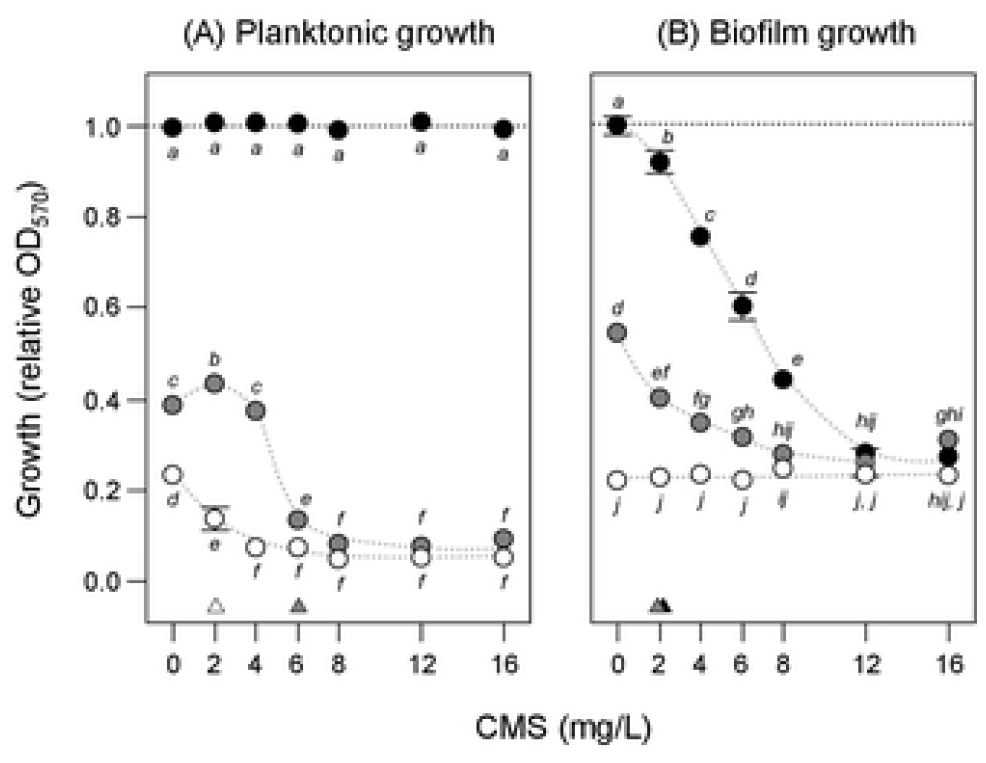
AZM and CMS synergistically inhibit *K. pneumonia* planktonic and biofilm growth. The effect of azithromycin (AZM) and colistin methanesulfonate (CMS) on *K. pneumonia* ATCC 10031 (A) planktonic and (B) biofilm growth was investigated by optical density (OD_570_) measurements after 16 h and 24 h incubation, respectively. Data are shown as mean relative OD_570_ ± SE (*n* = 3), with no-AZM (control) treatments indicated by dark circles, 3 mg/L AZM by grey circles and 9 mg/L AZM by white circles (trends are suggested by dashed curves). Means not connected by the same letters are significantly different (LSMeans Differences Tukey HSD, alpha = 0.05 for planktonic and biofilm growth mixed-effects models of relative OD_570_). Small triangles indicate planktonic and biofilm minimum inhibitory concentrations (MIC) where relative OD_570_ was first significantly reduced by CMS for each AZM treatment.

The effect of AZM and CMS on biofilm growth was further confirmed by MTT assays (**Fig. 4**) where a significant synergistic interaction was found in a mixed-effects model of relative A_570_ (*AZM x CMS, F*_12,12_ = 134, p = 0.01) as well as significant concentration effects (*AZM, F*_2,2_ = 2663, p = 0.01; *CMS, F*_6,6_ = 299, p = 0.01; data limited to 0 – 16 mg/L CMS to be consistent with the relative OD_570_ models). Again, relative A_570_ and OD_570_ treatment means were positively correlated (rho = 0.82, p = 0.01), demonstrating that the optical density and metabolic measurements of biofilm growth were in agreement. Finally, the combined AZM and CMS treatment was found to inhibit further growth of established biofilms by 0.8x (LSMeans Differences Tukey HSD, alpha = 0.05; see **S2 Fig**. for effect of combined AZM and CMS treatment on further ATCC 10031 biofilm growth) and a significant AZM effect and synergistic interaction were found in a mixed-effects model of relative A_570_ (*AZM, F*_1,1_ = 362, p = 0.01; *AZM x CMS, F*_2,2_ = 74, p = 0.01) but not a CMS effect (*CMS, F*_2,2_ = 0.2; p = 0.79). Our initial expectations were that CMS would have little effect on either planktonic or biofilm growth of *K. pneumonia* and that AZM is not considered an appropriate treatment for such infections. However, collectively our findings suggest that AZM and CMS have a synergistic activity on the planktonic growth of ATCC 10031 cells as well as on biofilm–formation. This suggests that an AZM-CMS combination therapy might be effective in treating biofilm–based Klebsiella hospital infections.

**Figure 4.**
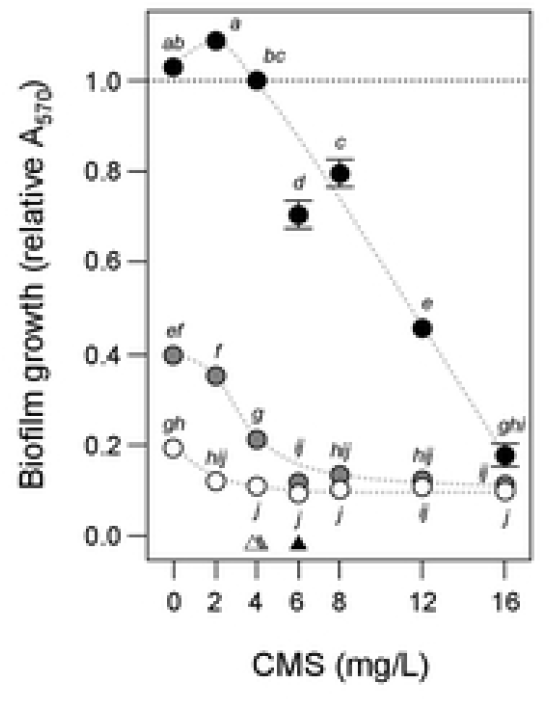
MTT assays of *K. pneumonia* biofilm growth confirms synergistic action of AZM and CMS. The effect of azithromycin (AZM) and colistin methanesulfonate (CMS) on *K. pneumonia* ATCC 10031 biofilm growth was investigated using the metabolic indicator dye MTT with absorbance (A_570_) measurements after 24 h incubation. Mean relative A_570_ ± SE (*n* = 3) is shown, with no-AZM (control) treatments indicated by dark circles, 3 mg/L AZM by grey circles and 9 mg/L AZM by white circles (trends are suggested by dashed curves). Means not connected by the same letters are significantly different (LSMeans Differences Tukey HSD, alpha = 0.05 for mixed-effects model of relative A_570_). Small triangles indicate planktonic and biofilm minimum inhibitory concentrations (MIC) where relative A_570_ was first significantly reduced by CMS for each AZM treatment.

### AZM and CMS in combination inhibits biofilm growth of *K. pneumonia* hospital isolates

As there are often significant differences between potentially laboratory-adapted model strains and recent hospital isolates, we investigated whether the synergistic action of AZM and CMS inhibiting the biofilm growth of *K. pneumonia* ATCC 10031 might also occur with hospital isolates. To investigate this, we chose five multiple drug resistant (MDR) and six non-MDR *K. pneumonia* Ukrainian hospital isolate (UHI) strains as *K. pneumonia* is the leading nosocomial gram-negative pathogen in Ukraine [1,2], and all of which were found to produce S-L interface biofilms under the conditions used here. The origins of the UHI strains as well as their antibiotic sensitivities are listed in **S1 Table** (note that although UHI 1090 was found to be sensitive to polymyxin B in the disc-diffusion assay, it was resistant to COL in the broth dilution assay whereas UHI 1633 was resistant in both assays). We then tested the inhibitory effect of AZM on the biofilm growth of these strains. Hierarchical cluster analysis based on antibiotic susceptibly clearly grouped these strains by MDR/non-MDR type (see **S3 Fig**. for the HCA dendrogram). Significant inhibition of biofilm growth was seen for four MDR strains and all non-MDR strains (LSMeans Differences Tukey HSD, alpha = 0.05 for individual mixed-effects models of OD_570_; see **S4 Fig**. for the effect of AZM on UHI strain biofilm growth). A combined mixed-effects model of OD_570_ with all strains confirmed significant strain and AZM effects (*MDR/non-MDR type, F*_1,1_ = 12, p = 0.01; *strain[type], F*_9,9_ = 26, p = 0.01; *AZM, F*_2,2_ = 221, p = 0.01). Despite the MDR/non-MDR division seen earlier, a hierarchical cluster analysis using these OD_570_ treatment means was found to cluster strains independently of strain type (Chi-square, DF_4,2_ = 4.95, p = 0.18; see **S3 Fig**. for the HCA dendrogram). The differences seen in biofilm response to AZM indicates that this small collection of strains was diverse and suggests that it could be used to test whether the synergistic action of AZM and CMS occurs more widely amongst *K. pneumonia* hospital isolates.

The effect on relative biofilm growth (OD_570_) of the MDR and non-MDR UHI strains by CMS in combination with AZM was then investigated (**Fig. 5**). A mixed-effects model of relative OD_570_ found significant strain effects (MDR & non-MDR type, *F*_1,1_ = 53, p = 0.3; strain[type], *F*_9,9_ = 28, p = 0.01) as well as a synergistic interaction across strains (AZM *x* CMS, *F*_2,2_ = 7, p = 0.01). Individual strain mixed-effects models of relative OD_570_ confirmed the synergistic interaction in all but UHI 1633 and 509 (UHI 1633, *F*_2,2_ = 2, p = 0.20; UHI 509, *F*_2,2_ = 0.4, p = 0.70) where it appears that the effects of AZM and CMS are simply additive. This test of UHI strains strongly suggests that the synergistic action of AZM and CMS is likely to be seen for other MDR and non-MDR *K. pneumonia* hospital isolates.

**Figure 5.**
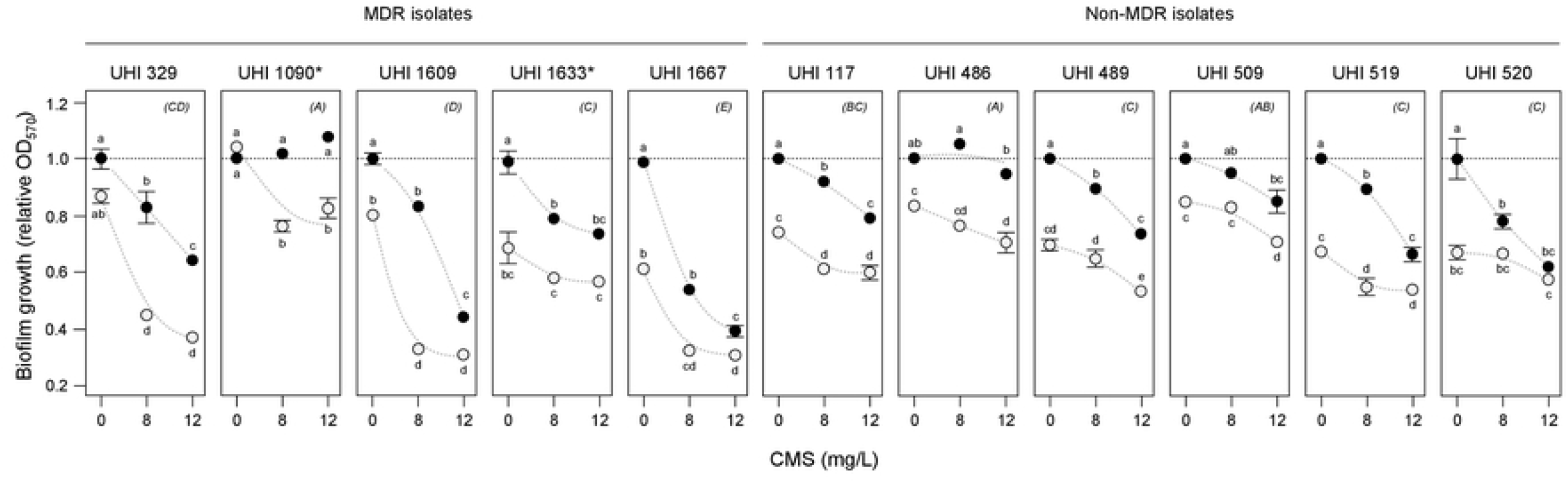
AZM and CMS synergistically inhibit biofilm growth of *K. pneumonia* hospital strains. The effect of azithromycin (AZM) and colistin methanesulfonate (CMS) on biofilm growth of multiple-drug resistant (MDR) and non-MDR *K. pneumonia* Ukrainian Hospital Isolate (UHI) strains was investigated using biofilm assays and with optical density (OD_570_) measurements after 24 h incubation. Mean relative OD_570_ ± SE (*n* = 3) is shown, with no-AZM (control) treatments indicated by dark circles and 9 mg/L AZM treatments by light circles, (trends are suggested by dashed curves). Means not connected by the same letters are significantly different (LSMeans Differences Tukey HSD, alpha = 0.05 for individual UHI strain mixed-effects models of relative OD_570_). Strains not connected by the same letters indicated in parentheses in the top-right of each panel are significantly different (LSMeans Differences Tukey HSD, alpha = 0.05 for combined UHI strain mixed-effects model of relative OD_570_). *, UHI 1090 and UHI 1633 were found to be resistant to COL in the broth dilution assay.

### Polymyxin B disc-diffusion inhibition of colony growth of *K. pneumonia* hospital isolates on AZM plates also shows synergistic action

We were interested in determining whether the synergistic action of AZM and CMS was also evident in antibiotic disc-diffusion assays which involve the inhibition of colony growth rather than biofilms. As CMS is not readily available in discs, we used the related antibiotic polymyxin B which is the preferred alternative for antimicrobial susceptibility testing when CMS is used in treatments (EUCAST recommendations [45]) (Polymyxin B and CMS (polymyxin E) are largely equivalent with the main differences associated with formulation and administration [46]). Although others have used Epsilometer test (E-test) strips [15] to deliver the first antibiotic over a range of concentrations onto an agar plate containing the second antibiotic to test for synergistic action (e.g., [47]), we found it easier and more accurate to measure colony inhibition using single-concentration discs placed on a range of plates with different concentrations of the second antibiotic. We measured the zone of colony inhibition (mm) surrounding polymyxin B discs on 0, 3, 6, 9 mg/L AZM plates spread with MDR and non-MDR *K. pneumonia* hospital isolates (AZM did not prevent colony growth at any concentration; **Fig. 6**). Significant increases in sensitivity to polymyxin B were seen with increasing AZM levels for most strains except UHI 329 and UHI 1663 (LSMeans Differences Tukey HSD, alpha = 0.05 for individual strain mixed-effects models but see **Fig. 6** legend for further details) and the synergistic activity confirmed by positive correlations between relative inhibition and AZM concentration for these strains (rho = 0.64 – 0.96, p = 0.01 – 0.03). The synergistic activity was the highest for UHI 509 with the inhibition zone diameter around the polymyxin B discs increasing 1.8x on 9 mg/L AZM plates, with the majority of strains only showing a 1.2x increase, and UHI 329, 1090, 1663 and 1667 showing little or no change (UHI 329, 1090 and 1667 had previously been shown to be sensitive polymyxin B while UHI 1663 was resistant, **S1 Table**; in this case there was no inhibition of colony growth next to the polymyxin discs on any of the AZM plates). Finally, a mixed-effects model of relative inhibition confirmed strain effects (*MDR & non-MDR type, F*_1,1_ = 27, p = 0.01; *strain[type], F*_8,8_ = 13, p = 0.01) in agreement with the earlier finding that there is diversity within this collection of MDR and non-MDR UHI hospital isolates, as well as an AZM effect across strains (*AZM, F*_3,3_ = 85, p = 0.01).

**Figure 6.**
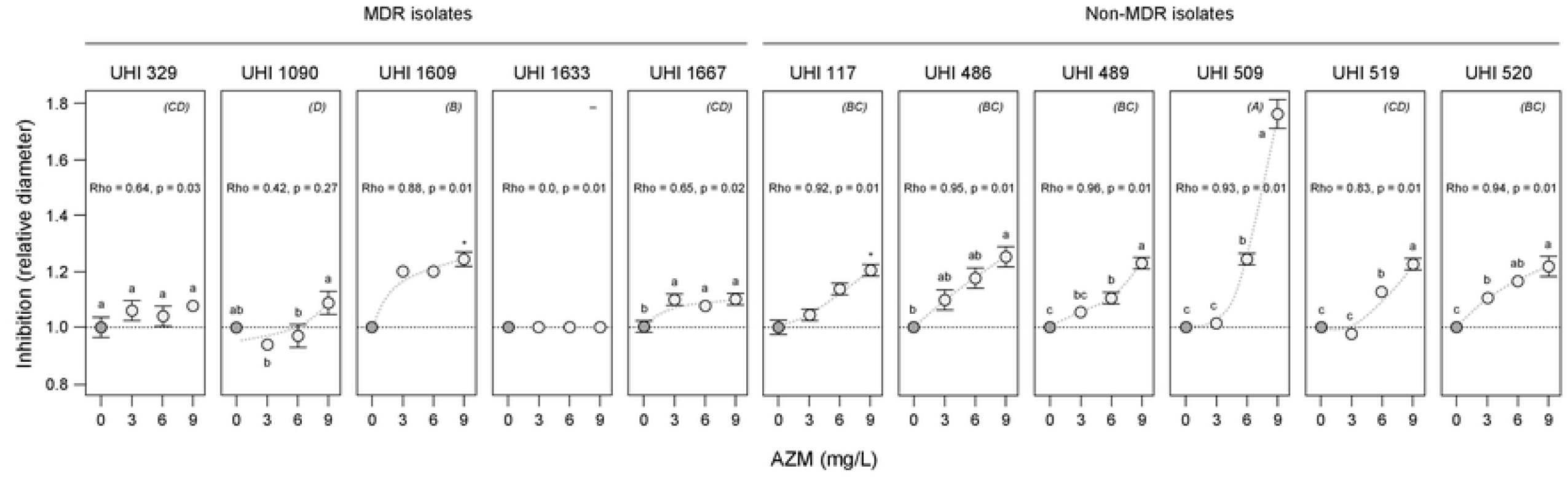
Synergistic inhibition of colony growth of *K. pneumonia* hospital strains by AZM and polymyxin B. The inhibition of colony growth of multiple-drug resistant (MDR) and non-MDR *K. pneumonia* Ukrainian Hospital Isolate (UHI) strains was investigated using polymyxin B disks on azithromycin (AZM) plates with colony inhibition zone measurements after 48 h incubation. Mean relative inhibition (mm) ± SE (*n* = 3) is shown, with no-AZM (control) treatments indicated by dark circles and 3 – 9 mg/L AZM treatments by light circles (trends are suggested by dashed curves). Means not connected by the same letters within panels are significantly different (LSMeans Differences Tukey HSD, alpha = 0.05 for individual mixed-effects models of relative inhibition for UHI 329, 509, 520, 1090 & 1667; individual general linear models for UHI 486, 489, 509, 519 & 1633; asterisk indicates significant difference between no AZM and 9 mg/L AZM treatments; Wilcoxon, UHI 117, ChiSquare, DF_1_ = 4.1, p = 0.04; UHI 1609, ChiSquare DF_1_ = 4.5, p = 0.03). Spearman’s correlation (Rho) between relative inhibition and AZM concentration is provided for each strain. Strains not connected by the same letters indicated in parentheses in the top-right of each panel are significantly different (LSMeans Differences Tukey HSD, alpha = 0.05 for combined UHI strain mixed-effects model of relative inhibition; UHI 1633 was omitted from this analysis as there was no variation in relative inhibition across AZM concentrations and was resistant to polymyxin B).

## Conclusions

The development of combination therapies based on the synergistic activities between antibiotics to overcome resistance is of growing interest to combat GNB pathogens such as *Klebsiella pneumoniae* [8,13]. This has involved the exploration of unusual combinations contraindicated by current treatment guidelines, and further opportunities in developing effective therapies may also be found in recognising that susceptibility testing and efficacy in patient treatments may be sensitive to bacterial lifestyles, as free-swimming planktonic and sessile biofilm-encased cells are known to have different physiology and resistance to stress and antibiotics. For example, most susceptibility testing in hospitals is based on the Kirby-Bauer disc-diffusion assay (sometimes using E-test strips) [14,15] in which colony (lawn) development is inhibited, but it is unclear whether this more closely resembles planktonic or biofilm growth (the disc diffusion assays are often followed by broth-dilution tests [15]). Furthermore, bacteria are known to grow in a continuum of aggregations which range from a few closely associated cells to highly densely packed structures in liquids, viscous environments and on surfaces [31], which suggests that bacterial growth is more nuanced, and the term ‘biofilm’ is often used without sufficient definition [31,48]. We have found that biofilm growth by a model *Klebsiella pneumoniae* strain (ATCC 10031) as well as hospital isolates can be inhibited by several macrolides including azithromycin (AZM), and that AZM can improve the effectiveness of colistin methanesulfonate (CMS), currently the last-agent-of-choice for *Klebsiella* infections [8]. However, in our investigation of ATCC 10031 and both MDR and non-MDR hospital isolates, we have found discrepancies suggesting that disc-diffusion and broth growth-based assays may not be good predictors of antibiotic susceptibility in biofilms. Better choices of combination therapies might be made if simple biofilm assays were used when determining which antibiotics to use in patient treatment.

## Acknowledgments

This work was undertaken with the support of the National Research Foundation of Ukraine through the Leading and Young Scientists Research Support programme and the research grant ‘Development of combined therapy for severe *Klebsiella pneumoniae*-associated nosocomial infections to overcome the antibiotic resistance’ (2020.02/0246). OVM and OSI also acknowledges Kyiv National University of Technologies and Design research grant ‘Development of a complex preparation of combined action based on collagen derivatives for the treatment of wound surfaces’ (0120U101290) and NATO SPS grant ‘Fighting maritime corrosion and biofouling with task-specific ionic compounds’ (984834). OMV thanks Viktoria Savchenko and the IMD Medical Laboratory for their support, and Galyna Filonenko, Centre for Paediatric Cardiology and Cardiac Surgery, Kiev, and Olga Balko, Zabolotny Institute of Microbiology and Virology of the National Academy of Sciences of Ukraine, for their technical support. OVM also thanks I. Gilbert and D. Purves, Wellcome Centre for Anti-Infectives Research, University of Dundee, for support to participate in the ‘Setting our sights on infectious diseases’ conference in 2019, and to L. Silver, LL Silver Consulting, New Jersey, USA, for productive discussions.

## Supporting information

**S1 Data Sets. Experimental data sets**. This file lists the quantitative replicate data used in the statistical analyses and figures presented in this work.

**S1 Figure. Macrolides differentially effect *K. pneumonia* biofilm growth**. Macrolides show different levels of inhibition of *K. pneumonia* ATCC 10031 biofilm growth and can be visualised by Hierarchical cluster analysis that group strains according to similarity. Dendrograms were constructed using mean relative (A) optical density (OD_570_) and (B) metabolic activity (A_570_) measurements of biofilm growth after 24 h. Macrolides linked to the same node (small circles) are more similar than those linked by deeper nodes, and the arbitrary root for this dendrogram is indicated by the dashed circle.

**S2 Figure. Combined AZM and CMS treatment supresses further *K. pneumonia* biofilm growth**. The effect of azithromycin (AZM) and colistin methanesulfonate (CSM) on established *K. pneumonia* ATCC 10031 biofilms was investigated by optical density (OD_570_) measurements after 24 h incubation. Mean relative OD_570_ ± SE (*n* = 3) is shown, with no-AZM (control) treatments indicated by dark circles and 9 mg/L AZM treatments by light circles (trends are suggested by dashed curves). Means not connected by the same letters are significantly different (LSMeans Differences Tukey HSD, alpha = 0.05 for mixed-effects model of relative OD_570_).

**S3 Figure. *K. pneumonia* hospital strains are differentiated by antibiotic susceptibility and AZM inhibition of biofilm growth**. The phenotypic diversity among the multiple-drug resistant (MDR) and non-MDR Ukrainian Hospital Isolate (UHI) strains can be visualised by Hierarchical cluster analysis that group strains according to similarity of (A) antibiotic susceptibility and (B) the effect of azithromycin (AZM) on biofilm growth in dendrograms with different topologies. The dendrogram on the left was constructed using disc-diffusion assay data for those antibiotics listed in Table S1 and the one on the right using mean relative optical density (OD_570_) measurements of biofilm growth with AZM after 24 h. MDR strains are indicated by dark circles and non-MDR strains by light circles. Strains linked to the same node (small circles) are more similar than those linked by deeper nodes, and the arbitrary roots for these dendrograms are indicated by the dashed circle.

**S4 Figure. AZM inhibits biofilm growth of *K. pneumonia* hospital strains**. The effect of azithromycin (AZM) on biofilm growth of multiple-drug resistant (MDR) and non-MDR *K. pneumonia* Ukrainian Hospital Isolate (UHI) strains was investigated by biofilm assays with optical density (OD_570_) measurements after 24 h incubation. Data are shown as mean OD_570_ ± SE (*n* = 3) (trends are suggested by dashed curves). Means not connected by the same letters are significantly different (LSMeans Differences Tukey HSD, alpha = 0.05 for individual UHI strain mixed-effects models of OD_570_; no significant effect for UHI 329, p = 0.06). Strains not connected by the same letters indicated in parentheses in the top-left of each panel are significantly different (LSMeans Differences Tukey HSD, alpha = 0.05 for combined UHIO strains mixed-effects model of relative inhibition).

**S1 Table. Origin and antibiotic susceptibility of the UHI *Klebsiella pneumoniae* strains used in this work**. This table lists the origins and antibiotic susceptibility of the UHI strains used in this work.

## Notes

### Competing Interest Statement

The authors have declared no competing interest.

